# Activity-regulated cytoskeleton-associated protein (Arc/Arg3.1)-mediated plasticity in the paraventricular thalamic nucleus promotes a fundamental adaptation to stress

**DOI:** 10.1101/2021.09.10.459826

**Authors:** Brian F. Corbett, Sandra Luz, Jay Arner, Abigail Vigderman, Kimberly Urban, Seema Bhatnagar

## Abstract

**BACKGROUND:** Habituation is defined as a progressive decline in response to repeated exposure to a familiar and predictable stimulus and is highly conserved across species. Disrupted habituation is a signature of post-traumatic stress disorder (PTSD). In rodents, habituation is observed in neural, neuroendocrine and behavioral responses to repeated exposure to the predictable and moderately intense stress or restraint. We previously demonstrated that lesions to the posterior division of the paraventricular thalamic nucleus (pPVT) impairs habituation. However, the underlying molecular mechanisms and specific neural connections among the pPVT and other brain regions that underlie habituation are unknown.

**METHODS:** Behavioral and neuroendocrine habituation was assessed in adult male Sprague-Dawley restraints using the repeated restraint paradigm. Pan neuronal and Cre-dependent Designer Receptors Exclusively Activated by Designer Drugs (DREADDs) were used to chemogenetically inhibit the pPVT and the subpopulation of pPVT neurons that project to the medial prefrontal cortex (mPFC), respectively. Activity-regulated cytoskeleton-associated protein (Arc) expression was knocked down in the pPVT using siRNA directed towards Arc. Golgi staining was used to assess structural plasticity of pPVT neurons. Local field potential recordings were used to assess coherent neural activity between the pPVT and mPFC. The attentional set-shifting task was used to assess mPFC-dependent behavior.

**RESULTS:** Here, we show that Arc promotes habituation by increasing stress-induced spinogenesis in the pPVT, increasing coherent neural activity with the mPFC, and improving mPFC-mediated cognitive flexibility.

**CONCLUSION:** Our results demonstrate that Arc induction in the pPVT regulates habituation to repeated restraint and mPFC function.

**One Sentence Summary:** We demonstrate that Arc in the posterior division of the paraventricular thalamic nucleus promotes habituation to repeated stress by increasing dendritic spines.

## Background

Habituation is defined as progressively reduced responsiveness to repeated exposure to the same predictable, mild to moderately intense stimulus (1-3). Habituation is displayed by species ranging from rodents to humans and is characterized in rodents by progressive reductions in behaviors and hypothalamic-pituitary-adrenal (HPA) axis responses to repeated experience with predictable and familiar homotypic stressors (1, 2, 4-6). Habituation is disrupted in individuals with post-traumatic stress disorder (PTSD) and is a key contributor to hyperarousal and re-experiencing symptom clusters (7-10). Additionally, habituation is critical for the success of behavioral therapies for PTSD (8, 11, 12). Habituation to stress is considered adaptive because it allows humans and animals to filter out irrelevant stimuli and focus selectively on important stimuli (13). However, a clear understanding of the molecular substrates underlying habituation to repeated stress has not emerged.

We have previously demonstrated that activity in the posterior division of the paraventricular thalamic nucleus (pPVT) is necessary for habituation (14). The PVT is an extensive midline thalamic nucleus divided into anterior, middle and posterior divisions based on segregation of afferent and efferent projections (15-20). Efferents of the anterior and medial divisions of the PVT are widespread (17, 21-23). In contrast, efferent projections of the pPVT are limited to: the central, basomedial, basolateral amygdala, nucleus accumbens, anterior olfactory nucleus, bed nucleus of the stria terminalis (BNST), peri-posterior paraventricular nucleus of the hypothalamus (peri-PVN), but not the PVN, and the medial prefrontal cortex (mPFC) (17, 18, 20, 24-27). The specific molecular substrates within the pPVT underlying habituation are not clear.

Activity-regulated cytoskeleton-associated protein (Arc/Arg3.1) is an immediate early gene that couples changes in neuronal activity with synaptic plasticity (28-30). Arc expression is increased within 30min of an activity-inducing event and returns to baseline within 4 hours (31). Arc is necessary for late-phase long-term potentiation (LTP) in the hippocampus (30, 32, 33). This is attributed to Arc’s role in interacting with proteins that stabilize and promote the branching of actin filaments (29, 32, 34, 35), thereby increasing dendritic spine numbers (36, 37). Indeed, Arc is required for hippocampus-dependent memory (30). Therefore, Arc-mediated mechanisms that have been identified in the hippocampus may also contribute to habituation to repeated stress by promoting neuronal plasticity in the pPVT.

Here, we show that Arc in the pPVT is induced following restraint, and that this induction is necessary for habituation to repeated restraint. We demonstrate that Arc promotes habituation through the formation of new spines on pPVT dendrites. Further, Arc in the pPVT regulates the coherence of oscillatory activity between the pPVT and the mPFC and mediates PFC-mediated cognitive flexibility in habituated animals. Together, these findings suggest that Arc-mediated synaptic plasticity in the pPVT and its regulation of projections to the mPFC underlie a fundamental, phylogenetically-conserved adaptation to stress.

## METHODS AND MATERIALS

### Animals

Adult male Sprague–Dawley rats (225–250 g) were obtained from Charles River Laboratories. Rats were singly housed with food and water available *ad libitum*. All rats were randomly assigned to groups by a lab member that was not involved with experiment procedures or data analysis. Rats were euthanized by rapid decapitation and their brains were immediately snap-frozen in 2-methylbutane or prepared for Golgi staining (see supplemental information).

### Restraint

Rats were restrained daily in clear, plexiglass tubes. On day 1 or day 5, they were euthanized 60min following restraint onset (4, 38, 39). Blood samples from the tail vein were collected to assess plasma adrenocorticotropic hormone (ACTH) and corticosterone at 0, 15, 30, and 60min following 30min restraint onset on days 1 and 5 (14, 38, 40). Plasma ACTH and corticosterone were quantified using a radioimmunoassay kit from MP Biomedical (Orangeburg, NY, USA) (see supplemental information).

### Immunohistochemistry

The following primary antibodies were used: mouse anti-Arc (156003 1:200, Synaptic Systems), mouse anti-mCherry (IC51, 1:100, Novus Biologicals), and rabbit anti-HA (C29F4, 1:250, Cell Signaling). Alexa Fluor ® secondary antibodies were used at a concentration of 1:200 (see supplemental information).

### Stereotaxic Injections and drug administration

The following siRNA constructs were used: iAAV-scramble (iAAV01508, serotype 8) and iAAV-Arc (iAAV06494008, serotype 8). Non-Cre-dependent AAV8-hSyn-hM4D-HA-IRES-mCherry and Cre-dependent AAV8-hSyn-DIO-hM4D-HA-mCherry were purchased from the University of North Carolina Vector Core. CAV2-Cre was purchased from Institut de Génétique Moléculaire de Montpellier, University of Montpellier. Surgeries for Arc knockdown were performed between 7-10 days prior to restraint. Surgeries for hM4D and Cre expression were performed 28 days prior to restraint. For DREADDs experiments, rats were intraperitoneally administered clozapine-*N*-oxide (CNO, 2 mg/kg) or vehicle (4% DMSO in saline) 60min prior to each restraint. For MK-8931 injections, cannulae (Plastic One, C313I/SPC) were placed to target the pPVT. Dummy cannulae were removed and 1 µL of either vehicle (10% DMSO in saline) or MK-8931 (Verubecestat, 100 ng/1µL, S8564, Selleckchem) was injected in awake rats (see supplemental information).

### Local field potential recordings

Recording electrodes were placed in the mPFC and within 0.5 µM of the AAV injection site in the pPVT. Baseline (5min) and recovery (10min) recordings were performed in the home cage. Restraint recording (first 15min) were performed in a larger restraint tube (see supplemental information).

### Golgi Staining

Golgi staining was performed using the Rapid GolgiStain™ kit (FD Neurotechnologies, PK401) per manufacturer instructions. Neurons were traced at 100x using Neurolucida. In the pPVT and aPVT, typically 2 neurons in 4 sections were traced for each rat. Group sizes indicate the number of rats in each group (see supplemental information).

### Attentional Set Shifting Task

Following 5 consecutive days of restraint, iAAV-scramble and iAAV-Arc rats underwent the attentional set shifting task (AST) as in (40) to assess the role of pPVT Arc knockdown on PFC function following restraint. Rats were tested in side discrimination, side reversal, and light discrimination phases. The number of trials required to reach criterion (8 consecutive correct trials), time required to reach criterion, number of omissions, and number of errors during the test phase were recorded and analyzed using MATLAB (see supplemental information).

### Statistical Analyses

Student’s t-tests, one-way ANOVAs, ordinary two-way ANOVAs, and repeated measures two-way ANOVAs were performed in GraphPad Prism 7 (see supplemental information, figure legends, and supplemental Tables 1 and 2).

## RESULTS

### Habituation induces Arc expression in the pPVT but not in the anterior (aPVT)

Consistent with previous results (4, 14, 40), we confirmed that plasma ACTH and corticosterone concentrations were reduced during the 5^th^ day of restraint (Figure 1A,B and Supplemental Figure 1A,B). By the 3^rd^ restraint, struggle duration was significantly lower than on day 1 (Figure 1C). The density of Arc-immunoreactive (IR) neurons was increased in the pPVT (Figure 1D,E), but not the aPVT (Figure 1F), of rats restrained for 1 and 5 days compared to non-restrained controls. The density of c-Fos-IR cells is also increased in the pPVT, but not the aPVT, of rats restrained for 1 or 5 days compared to non-restrained controls (Supplemental Figure 1C-E). Together, these results confirm that restraint induces neuronal activity in the pPVT.

**Figure 1.**
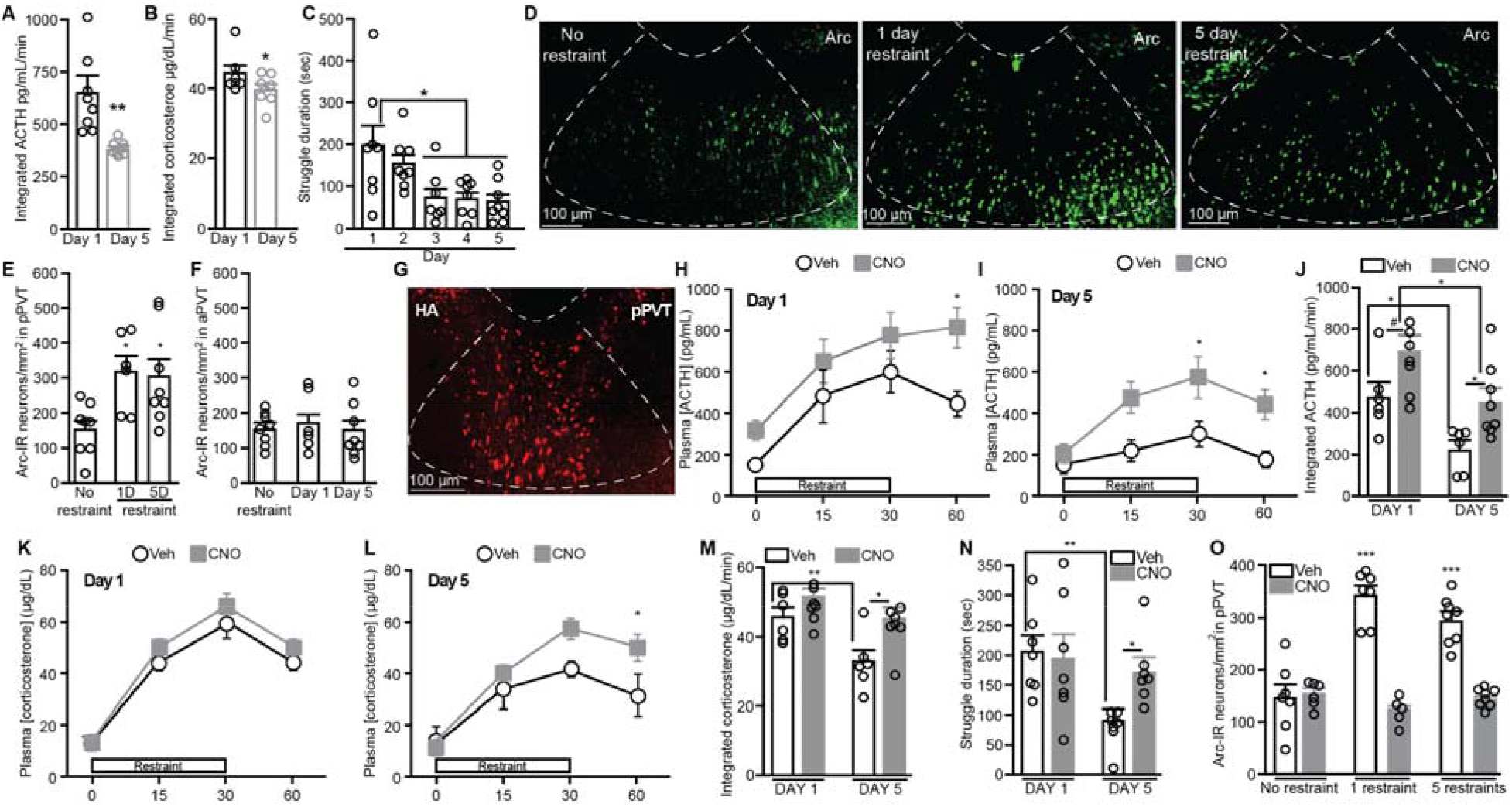
Chemogenetic inhibition of the pPVT impairs habituation to repeated restraint. **A)** Integrated plasma ACTH and **B)** corticosterone from 0-60min following restraint onset in naïve rats (*n* = 8/day). **C)** Struggle duration during the first 15min of restraint in naïve rats (*n* = 8/day). **D)** pPVT Images of and quantification of Arc-IR neurons in the **E)** pPVT and **F)** aPVT (no restraint *n* = 8, 1day *n* = 6, 5day *n* = 8). **G)** Image of HA tag in hM4D-expressing pPVT neurons. Plasma ACTH in vehicle-(*n* = 6) and CNO-treated rats (*n* = 8) during restraint on **H)** day1 and **I)** day5. **J)** Integrated plasma ACTH in vehicle-(*n* = 6) and CNO-treated rats (*n* = 8). Plasma corticosterone in vehicle-(*n* = 6) and CNO-treated rats (*n* = 8) during restraint on **K)** day1 and **L)** day5. **M)** Integrated plasma corticosterone in vehicle-(*n* = 6) and CNO-treated rats (*n* = 8). **N)** Struggle duration in vehicle- and CNO-treated rats (*n* = 7/group). **O)** Density of pPVT Arc-IR neurons in vehicle- and CNO-treated rats (no restraint vehicle *n* = 7, no restraint CNO *n* = 6, 1day vehicle *n* = 7, 1day CNO *n* = 5, 5day *n* = 8/treatment). Bars represent mean ± SEM. *p<0.05, **p<0.01, ***p<0.001, #p<0.10. **A**,**B**, paired Student’s t-test. **C**,**E**, Tukey’s multiple comparisons, one-way ANOVA. **E**, *different from controls. **H-N**, Sidak’s multiple comparisons, repeated measures two-way ANOVA. **O**, ***different from all groups except each other, Tukey’s multiple comparisons, ordinary two-way ANOVA.

### Chemogenetic inhibition of the pPVT impairs habituation

In otherwise naïve rats, CNO alone did not affect plasma concentrations of ACTH or corticosterone at any timepoint during days 1 or 5 of restraint (Supplemental Figure 2A-F) nor did CNO affect struggle duration (Supplemental Figure 2G). In rats expressing hM4D in the pPVT, CNO treatment increased plasma ACTH concentrations at the 60min timepoint on day 1 (Figure 1H) and the 30- and 60-min timepoints on day 5 (Figure 1I) compared to vehicle-treated controls. Integrated ACTH was also increased on days 1 and 5 in CNO-treated rats (Figure 1J). CNO treatment did not affect plasma corticosterone concentrations during day 1 of restraint (Figure 1K). However, compared to vehicle-treated controls, CNO-treated rats displayed increased plasma corticosterone concentrations 60min following restraint onset on day 5 of restraint (Figure 1L). Vehicle-treated, but not CNO-treated, rats displayed reduced integrated corticosterone on day 5 compared to day 1. CNO-treated rats displayed increased integrated plasma corticosterone concentrations compared to vehicle-treated controls on day 5 (Figure 1M). Vehicle-treated, but not CNO-treated, rats also displayed reduced struggle duration on day 5 compared to day 1. CNO-treated rats displayed increased struggle duration compared to vehicle-treated controls on day 5 (Figure 1N). Indeed, CNO treatment in hM4D-expressing rats chemogenetically inhibited the pPVT as vehicle-, but not CNO-treated, rats displayed increased densities of Arc-IR neurons in the pPVT 60min following the onset of 1 or 5 restraints (Figure 1O). Together, our findings indicate that chemogenetic inhibition of the pPVT impairs neuroendocrine and behavioral stress habituation.

### Stress-induced increases in Arc expression in the pPVT are necessary for habituation

We investigated whether Arc in the pPVT regulates habituation to repeated stress by using siRNA directed toward Arc (iAAV-Arc) to knockdown Arc expression. Arc-IR neuron density was significantly reduced in iAAV-Arc rats compared to iAAV-scramble controls following 5 consecutive daily restraints (Figure 2A,B), thereby confirming siRNA efficacy. In iAAV-scramble controls, ACTH concentrations were significantly reduced 15- and 30-min following restraint onset on day 5 compared to day 1 (Figure 2C). In iAAV-Arc rats, ACTH concentrations were similar at all timepoints on days 1 and 5 of restraint (Figure 2D). iAAV-scramble, but not iAAV-Arc, rats displayed reduced integrated ACTH on day 5 compared to day 1 of restraint (Figure 2E). In iAAV-scramble controls, corticosterone concentrations were significantly reduced 60min following restraint onset on day 5 compared to day 1 (Figure 2F). In iAAV-Arc rats, corticosterone concentrations were similar at all timepoints on days 1 and 5 of restraint (Figure 2G). iAAV-scramble rats displayed reduced integrated corticosterone on day 5 compared to day 1 of restraint but this decrease was not observed in iAAV-Arc rats (Figure 2H). iAAV-scramble rats displayed reduced struggle duration on day 5 of restraint compared to day 1 but iAAV-Arc rats did not show this reduction (Figure 2I). Together, these results indicate that restraint-induced Arc in the pPVT is necessary for behavioral and neuroendocrine habituation to repeated restraint stress.

**Figure 2.**
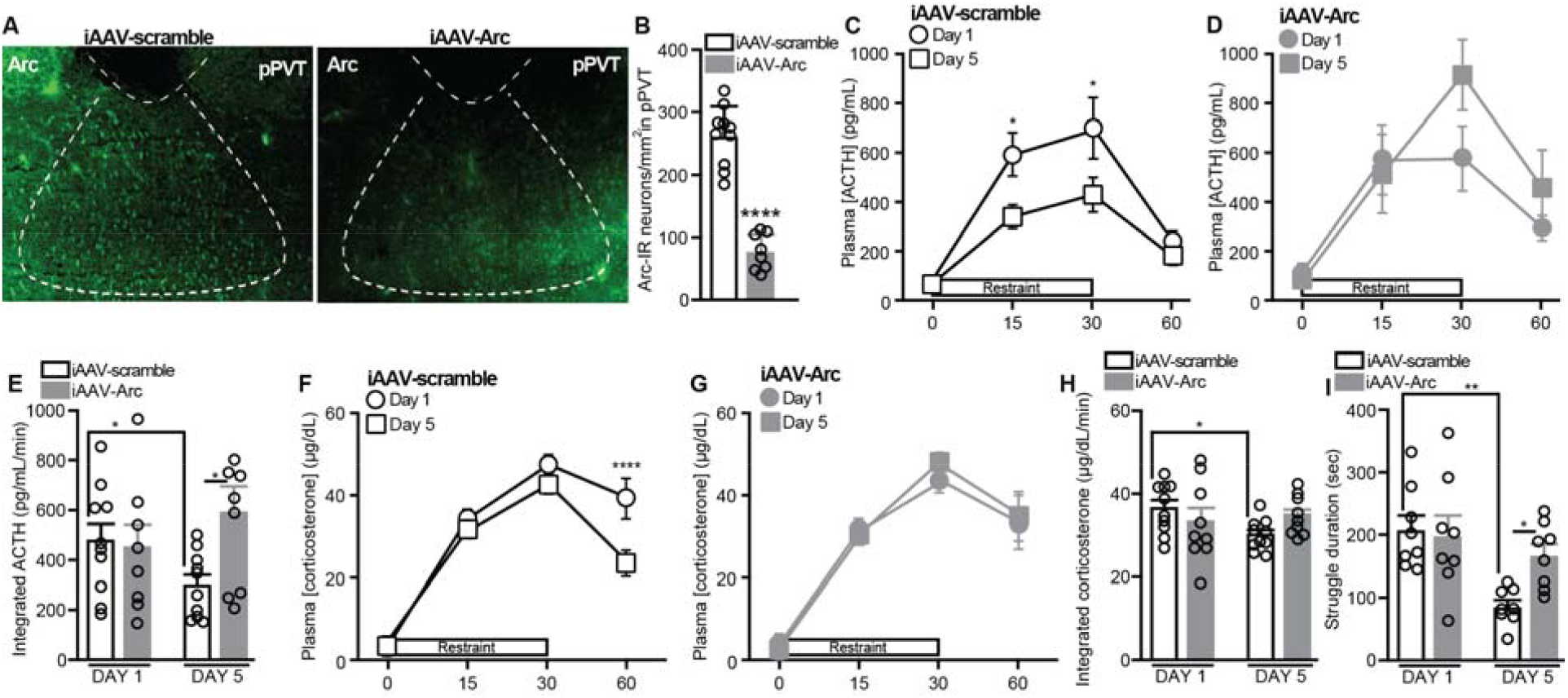
Arc in the pPVT regulates habituation to repeated restraint. **A)** Images of Arc in the pPVT of iAAV-scramble and iAAV-Arc rats. **B)** Density of Arc-IR neurons in the pPVT of iAAV-scramble (*n* = 10) and iAAV-Arc rats (*n* = 8) following 5 days of restraint. Plasma ACTH concentrations at 0, 15, 30, and 60min following the onset of 30min restraint on days 1 and 5 in **C)** iAAV-scramble (*n* = 10) and **D)** iAAV-Arc rats (*n* = 8). **E)** Integrated plasma ACTH concentrations in iAAV-scramble (*n* = 10) and iAAV-Arc rats (*n* = 10) from 0-60min following restraint onset on days 1 and 5. Plasma corticosterone concentrations at 0, 15, 30, and 60min following the onset of 30min restraint on days 1 and 5 in **F)** iAAV-scramble (*n* = 10) and **G)** iAAV-Arc rats (*n* = 8). **H)** Integrated plasma corticosterone concentrations in iAAV-scramble (*n* = 10) and iAAV-Arc (*n* = 8) rats from 0-60min following restraint onset on days 1 and 5. **I)** Struggle duration during the first 15min of restraint in iAAV-scramble and iAAV-Arc rats on days 1 and 5 (*n* = 8/group). Bars represent mean ± SEM. For **B**, ****p < 0.0001, Student’s unpaired t-test. For **C-I**, *p < 0.05, **p < 0.01, ***p < 0.001, Sidak’s multiple comparisons following repeated measures two-way ANOVA. Horizontal bars indicate differences between groups.

### Arc knockdown in the pPVT prevents stress-induced spinogenesis

We investigated whether Arc regulates structural plasticity in the pPVT. Arc knockdown had no effect on spine density in non-restrained rats (Figure 3A,B). 24 hours following a 5^th^ consecutive restraint, iAAV-Arc rats displayed reduced dendritic spine densities in the pPVT 80 and 100 µm distal from the soma (Figure 3C). Collapsing across distance from soma, mean pPVT spine density was increased by repeated restraint in iAAV-scramble, but not in iAAV-Arc rats, resulting in lower spine densities in iAAV-Arc rats compared to scramble controls on day 5 (Figure 3D). Mean dendrite length in the pPVT was not affected by restraint or Arc knockdown (Supplemental Figure 3A-C), indicating that effects on spine density cannot be solely attributed to changes in dendritic length. Together, these data indicate that Arc is necessary for restraint-induced increases in pPVT spine density.

**Figure 3.**
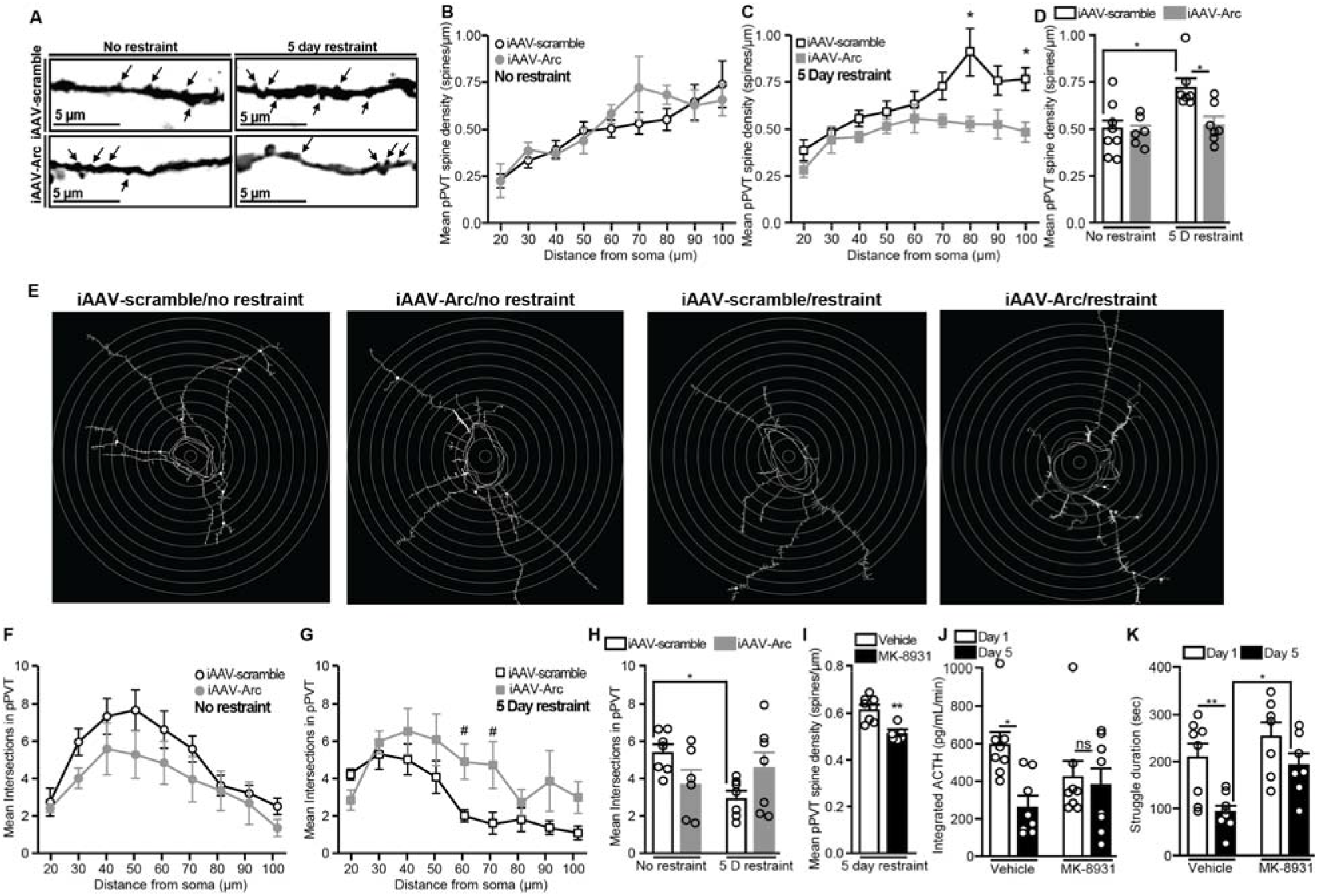
Arc knockdown prevents stress-induced spine densities and dendritic remodeling in the pPVT. **A)** Images of dendritic stubby spines in pPVT neurons. Mean spine density at increasing 10 µm radial increments in the pPVT of **B)** non-restrained controls (iAAV-scramble *n* = 8, iAAV-Arc *n* = 6) and **C)** 24 h following a 5^th^ daily restraint (*n* = 7/group). **D)** Mean pPVT spine density over all distances from the soma. **E)** Images of dendritic complexity in the pPVT. Mean number of intersections at increasing 10 µm radial increments from the soma in the pPVT of **F)** non-restrained controls (iAAV-scramble *n* = 7, iAAV-Arc *n* = 6) and **G)** 24 h following a 5^th^ daily restraint (*n* = 7/group). **H)** Mean dendritic intersections averaged over all distances from the soma (no restraint iAAV-scramble *n* = 7, no restraint iAAV-Arc *n* = 6, 5day restraint *n* = 7/group). **I)** Mean spine density of pPVT neurons averaged over all distances from the soma in vehicle-(*n* = 8) and MK-8931-treated rats (*n* = 7). **J)** Integrated plasma ACTH concentrations in vehicle- and MK-8931-treated rats from 0-60min following restraint onset (*n* = 7/group). **K)** Struggle duration during the first 15min of restraint in vehicle-(*n* = 8) and MK-8931-treated rats (*n* = 7). Bars represent mean ± SEM. *p < 0.05, **p < 0.01, #p < 0.10. **B, C, F, G, J**, and **K**, Sidak’s multiple comparisons, repeated measures two-way ANOVA. **D** and **H**, Tukey’s multiple comparisons, ordinary two-way ANOVA. **I**, Student’s t-test.

### Arc knockdown in the pPVT prevents reductions in dendritic complexity caused by stress

Arc knockdown did not affect dendritic complexity in pPVT neurons in the absence of stress as non-restrained iAAV-scramble and iAAV-Arc rats displayed similar numbers of dendritic intersections at all distances from the soma (Figure 3E,F). Following restraint, iAAV-Arc rats display trends for increased dendrite intersections 60 and 70 µm from the soma in pPVT neurons compared to iAAV-scramble controls (Figure 3G). Collapsing across distance from the soma, restraint reduced dendritic intersections in the pPVT of iAAV-scramble rats but not in iAAV-Arc rats (Figure 3H). Taken together, these findings indicate that stress reduces dendritic complexity in pPVT neurons and that this reduction is dependent on Arc. These structural effects are specific to pPVT neurons as aPVT neurons exhibited no restraint- or Arc-mediated effects on dendritic spine density (Supplemental Figure 3D-F), dendrite intersections (Supplemental Figure 3G-I), or dendrite length (Supplemental Figure 3J-L).

### Inhibition of spine formation in the pPVT impairs habituation

We next investigated whether the formation of new spines in the pPVT is necessary for habituation. We used the β-site amyloid precursor protein cleavage enzyme 1 (BACE1) inhibitor MK-8931, which inhibits the formation of new dendritic spines but not the stability of existing ones (41). The use of more common pharmacological inhibitors of spinogenesis, such as latrunculin A, could not be used as they inhibit actin dynamics (42) and therefore actin-mediated localization of Arc to the synapse (43, 44). This would be problematic as it would be impossible to determine whether spinogenesis or other Arc-mediated processes, such as reducing dendritic complexity, regulate habituation. Following restraint, total dendritic spines in the pPVT were reduced in MK-8931-treated rats compared to vehicle-treated controls (Figure 3I, Supplemental Figure 3M). MK-8931 treatment did not affect dendritic branching (Supplemental Figure 3N,O) or dendrite length (Supplemental Figure 3P,Q) in the pPVT. MK-8931 treatment impaired habituation as vehicle-treated, but not MK-8931-treated, rats exhibited reduced plasma ACTH concentrations (Figure 3J, Supplemental Figure 3R,S) and struggle duration (Figure 3K) on day 5 of restraint compared to day 1. These findings indicated that inhibiting restraint-induced spinogenesis in the pPVT is sufficient to impair habituation to repeated restraint.

### Arc knockdown in the pPVT reduces network activity in the pPVT and reduces coherent activity between the pPVT and mPFC in repeatedly stressed rats

The mPFC inhibits HPA activity (45, 46) and is the primary cortical target of the pPVT (26, 47-49). We hypothesized that Arc in the pPVT regulates habituation, at least in part, via projections to the mPFC. Power spectral density (PSD) was analyzed to assess network activity in the delta (2-4 Hz), theta (4-9 Hz), alpha (9-12 Hz), beta (12-20 Hz), and gamma (20-40 Hz) frequency bands before, during, and after restraint. Compared to iAAV-scramble rats, iAAV-Arc rats displayed reduced delta power in the pPVT during baseline and during the 5-10min bin following restraint on day 5 (Figure 4A, Supplemental Figure 4D). Compared to iAAV-scramble controls, iAAV-Arc rats also displayed reduced alpha power during the 5-10min bin of restraint on day 5 (Supplemental Figure 4E). Arc knockdown did not alter power in the pPVT in the theta, beta, or gamma frequency ranges at any timepoints (Supplemental Figure 4F-H). Arc knockdown in the pPVT did not alter power in any frequency ranges in the mPFC at any time points (Supplemental Figure 4I-N). Arc knockdown had no effect on pPVT-mPFC coherence during the day 1 baseline recording. During the day 5 baseline recording, iAAV-Arc rats displayed reduced pPVT-mPFC coherence in the delta (Figure 4C), theta (Figure 4D), and alpha (Figure 4E) frequency ranges compared to iAAV-scramble controls. During the 0-5min bin following day 1 restraint, iAAV-Arc rats displayed reduced pPVT-mPFC coherence in the delta range (Figure 4C) and a trend for reduced coherence in the theta range (Figure 4D) compared to iAAV-scramble controls. Following day 5 restraint, iAAV-Arc rats displayed reduced pPVT-mPFC coherence in the theta range 0-5min following restraint a trend for reduced theta coherence 5-10min following restraint. Arc knockdown in the pPVT did not affect pPVT-mPFC coherence at any time point in the beta or gamma frequency ranges (Supplemental Figure 4O,P). Together, these findings indicate that Arc expression in the pPVT regulates population activity in the pPVT important for delta and alpha frequency bands. Arc in the pPVT also promotes coherent neural activity between the pPVT and the mPFC during habituation.

**Figure 4.**
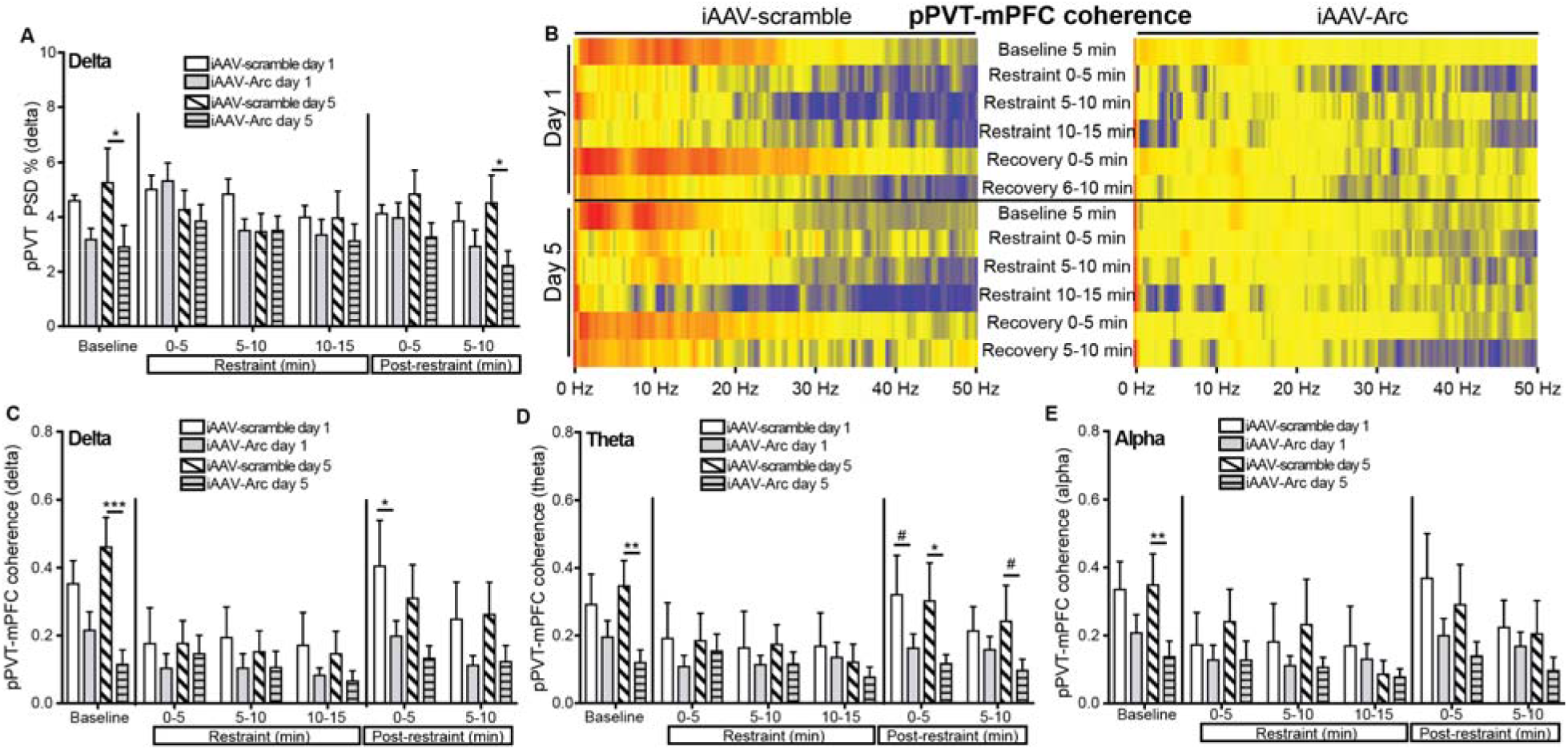
Arc knockdown in the pPVT disrupts pPVT-mPFC coherence. **A)** Delta PSD percentages in the pPVT of iAAV-scramble and iAAV-Arc rats during baseline, restraint, and recovery on days 1 and 5 of restraint (iAAV-scramble *n* = 6, iAAV-Arc *n* = 8). **B)** Images of mean pPVT-mPFC coherence value heat maps from 0-50 Hz in the pPVT of iAAV-scramble and iAAV-Arc rats during a 5 min baseline, three 5-min restraint bins, and two 5-min recovery bins on days 1 and 5 of restraint. Warmer colors represent higher coherence. Coherence values in the **C)** delta, **D)** theta, and **E)** alpha frequency ranges in the pPVT of iAAV-scramble and iAAV-Arc rats during baseline, restraint, and recovery on days 1 and 5 of restraint (iAAV-scramble *n* = 6, iAAV-Arc *n* = 8). Bars represent mean ± SEM. For **A** and **C-E, \***p < 0.05, **p < 0.01, ***p < 0.001, # p < 0.10, Fisher’s least significant difference following two-way ANOVA. Horizontal bars indicate differences between groups.

### Chemogenetic inhibition of mPFC-projecting pPVT neurons impairs behavioral but not neuroendocrine habituation

We next determined whether mPFC-projecting pPVT neurons impair habituation. CAV2-Cre, which transduces axon terminals to retrogradely express Cre recombinase, was injected into the mPFC. The pPVT was injected with AAV8-hSyn-DIO-hM4D-HA-mCherry, which expresses the inhibitory hM4D DREADD in a Cre-dependent manner (17). Sixty min following the onset of the 5^th^ restraint, the numbers of Arc-IR (Figure 5A,B) and c-Fos-IR (Supplemental Figure 5A,B) neurons in the pPVT that were also labeled with mCherry were reduced in CNO-treated rats compared to vehicle-treated controls. Vehicle-treated rats displayed reduced struggle duration during the 5^th^ restraint compared to the 1^st^ restraint but CNO-treated rats did not show this reduction (Figure 5C). These results indicate that mPFC-projecting pPVT neurons regulate behavioral habitation.

**Figure 5.**
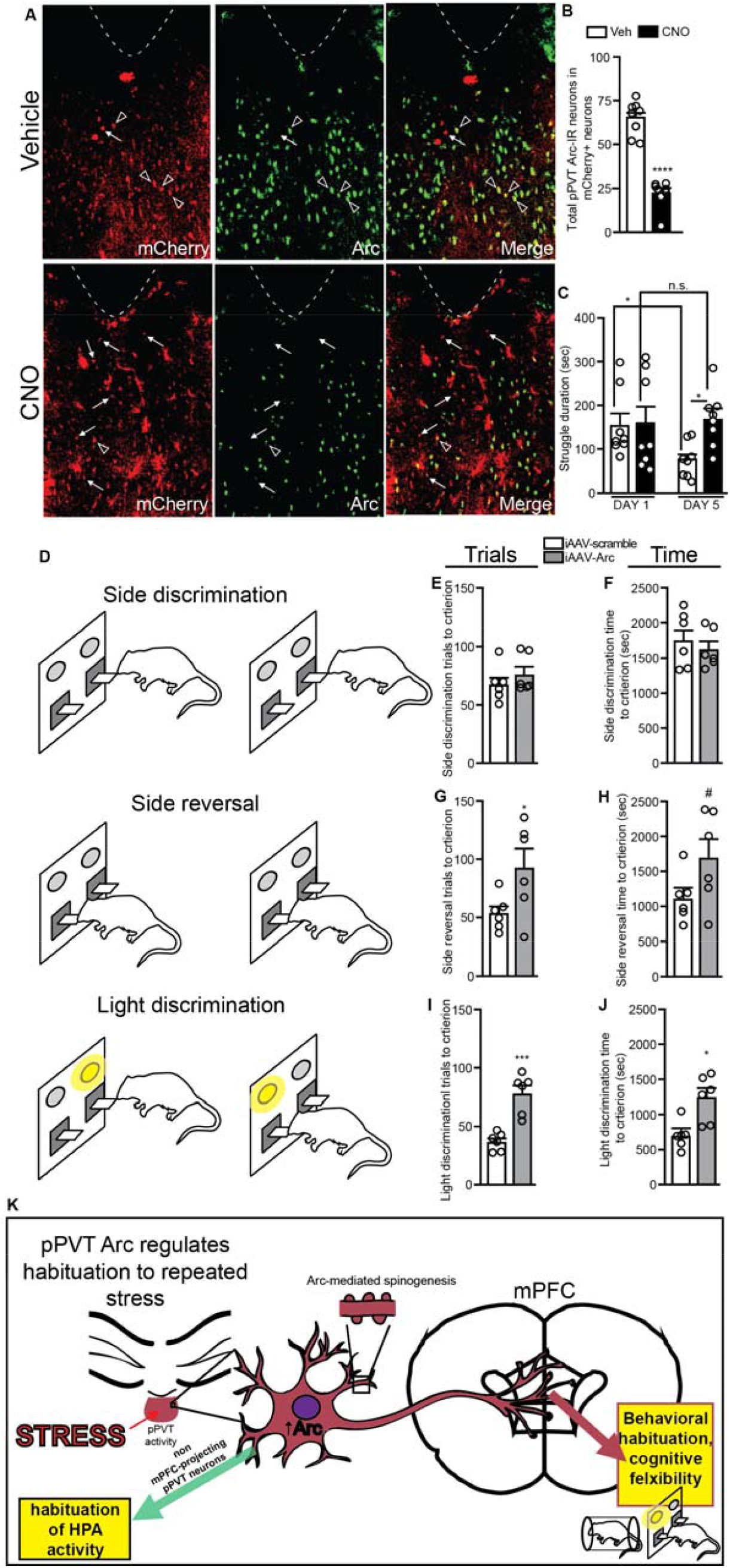
The pPVT projections to the mPFC regulate struggle habituation and mPFC function. **A)** Images of mCherry, Arc, and merged images from the pPVT of rats injected with CAV2-Cre in the mPFC and with AAV8-hSyn-DIO-hM4D-HA-mCherry in the pPVT after 5 restraints. **B)** The number of Arc+/mCherry+ neurons in the pPVT (*n* = 8/group). **C)** Struggle duration on days 1 and 5 of restraint vehicle- and CNO-treated rats (*n* = 8/group). **D)** Descriptive graphic of AST methods. The **E)** number of trials and **F)** time required to reach criterion (8 consecutive correct trials) during the side discrimination phase of the AST (*n* = 6/group). The **G)** number of trials and **H)** time required to reach criterion during the side reversal phase of the AST (*n* = 6/group). The **I)** number of trials and **J)** time required to reach criterion during the light discrimination phase of the AST (*n* = 6/group). **K)** Figure summarizing findings. Stress increases pPVT activity, which induces Arc. Arc-mediated increases in dendritic spine density facilitate habituation by sensitizing stress-activated input to pPVT neurons, particularly those projecting to the mPFC. pPVT Arc promotes pPVT-mPFC coherence and regulates behavioral habituation and cognitive flexibility. pPVT neurons that do not project to the mPFC may negatively regulate the HPA axis. Bars represent mean ± SEM. For **B** and **G-J**, *p < 0.05, ***p < 0.001, ****p < 0.0001, #p < 0.10, Student’s t-test. For **C**, *p < 0.05, Sidak’s multiple comparisons, repeated measures two-way ANOVA. Horizontal bars indicate differences between groups.

### Restraint-induced Arc in the pPVT promotes mPFC-dependent cognitive flexibility

We hypothesized that pPVT Arc knockdown would impair performance in the attentional set shifting task (AST) following restraint, particularly in the mPFC-dependent light discrimination phase (50), as Arc knockdown disrupted coherent activity with the mPFC. Arc knockdown did not affect the number of trials (Figure 5E) or time required to reach criterion (Figure 5F) in the side discrimination phase. In the side reversal phase, iAAV-Arc rats displayed an increased number of trials (Figure 5G) and a trend towards increased time (Figure 5H) required to reach criterion compared to iAAV-scramble rats. In the light discrimination phase, iAAV-Arc rats displayed an increased number of trials (Figure 5I) and increased time (Figure 5J) required to reach criterion compared to iAAV-scramble rats. Arc knockdown in the pPVT did not affect the percentage of omissions (Supplemental Figure 5C-E) or number of errors (Supplemental Figure 5F-H) in the AST. Together, these findings indicate that as male rats habituate to repeated restraint stress, induction of Arc in the pPVT contributes to improvements in subsequent mPFC-mediated cognitive flexibility.

## Discussion

Here, we demonstrate that neuronal activity in the pPVT produced by repeated stress induces the expression of Arc, which promotes habituation by increasing dendritic spine densities in the pPVT. Restraint-induced pPVT Arc is required for habituation since Arc knockdown in the pPVT impaired reductions in struggle behavior, ACTH, and corticosterone following multiple restraints. Arc knockdown prevented stress-induced increases in spine density and reduced dendritic complexity in pPVT neurons. Thus, Arc mediates stress-induced reformatting of pPVT neurons. Stress-induced spinogenesis contributes to habituation as pharmacological inhibition of spine formation in the pPVT using MK-8931 was sufficient to impair habituation. Increased spine density in the pPVT may sensitize pPVT neurons to be more responsive to specific afferents activated by restraint. Arc-mediated increases in pPVT spine densities may increase activity of projections to the mPFC as pPVT Arc knockdown reduced pPVT-mPFC coherence. We hypothesize that pPVT-mPFC coherence is an important mechanism underlying habituation as pPVT Arc knockdown impaired PFC-dependent cognitive flexibility and chemogenetic inhibition of mPFC-projecting pPVT neurons impaired behavioral habituation. These findings are important as they are, to best of our knowledge, the first to identify 1) a mechanism of neuronal plasticity in the pPVT that promotes a fundamental adaptation to repeated stress and 2) that restraint-induced pPVT Arc induction is important for cortically-dependent cognitive flexibility. Together, these findings offer novel insights into the molecular, cellular, and network mechanisms through which individuals reduce their responsivity to familiar and predictable stimuli and subsequently show improved cognitive flexibility.

While restraint-induced expression of other immediate early genes in the pPVT may also contribute to habituation, we focused on Arc because of its well-established role in regulating structural plasticity (29, 51, 52), a property of neurons that is also tightly regulated by stress (53, 54). We found that Arc is necessary for restraint-induced spinogenesis in the pPVT and that spinogenesis is necessary for habituation as inhibiting spinogenesis with MK-8931 impaired habituation. In addition to increasing spine density, restraint-induced Arc also mediated reductions in dendritic branching of pPVT neurons. Stress-mediated reductions in dendritic branching have been described in stress-related brain regions including the pPVT, mPFC, and hippocampus (53, 55, 56). In the hippocampus, NMDA receptors are necessary for reductions in the dendritic complexity caused by stress (57). Arc is a key link between NMDA receptor activity and structural remodeling. Therefore, Arc contributes to stress-mediated reductions in dendritic complexity in the pPVT and perhaps in other regions as well.

We report that in the pPVT of iAAV-Arc rats, power in the delta frequency range is reduced during the baseline recording and during recovery following restraint on day 5 compared to iAAV-scramble controls. Reduced delta power may reflect a decrease in quiet wakefulness (58-61) resulting from anticipatory activity prior to restraint and impaired return to quiet wakefulness following restraint, respectively. Power in the alpha range was also reduced during restraint on day 5 in iAAV-Arc rats. This may correlate with impairments in information processing and the filtering of sensory stimuli required for optimal attention (62, 63). Attention is essential for establishing restraint stress as not being physically threatening, a key aspect that makes habituation to these stressors adaptive (2). Together, reduced delta and alpha power in the pPVT of iAAV-Arc rats suggests a heightened state of arousal that may impair the ability to attend to restraint stress.

We hypothesized that pPVT projections to the mPFC regulate habituation. The mPFC is the primary cortical target of the pPVT (23, 26, 48) and inhibits activity in key stress-related brain regions including the amygdala (64, 65) and inhibits HPA axis activity (45, 46). pPVT-mPFC coherence was reduced in the delta, theta, and alpha frequency ranges during certain baseline and/or recovery bins in iAAV-Arc rats compared to iAAV-scramble controls. These reductions in coherence suggest that functional connectivity between the pPVT and mPFC may be impaired as the PVT directly projects to the mPFC (26, 47-49). It is worth noting that changes in coherence may not necessarily be caused by impairments in mPFC-projecting pPVT neurons as other structures, such as the locus coeruleus (LC), may modulate both pPVT (66) and mPFC activity in parallel (67). Regardless, reductions in synchronous oscillations between the pPVT and mPFC may impair the ability of the pPVT to influence mPFC activity. To directly examine whether pPVT-mPFC projections regulate habituation, we demonstrated that inhibition of mPFC-projecting pPVT neurons impaired behavioral habituation. Together, these findings demonstrate that pPVT Arc promotes pPVT-mPFC coherence and that mPFC-projecting pPVT neurons regulate behavioral habituation during repeated restraint.

We then examined whether Arc in the pPVT regulates mPFC-mediated behaviors. In previous work, we showed improved cognitive flexibility in the AST, a PFC-dependent task, in male rats that have habituated compared to non-stressed controls (40). Because Arc in the pPVT is increased by repeated restraint and contributes to pPVT-mPFC coherence, we hypothesized that pPVT Arc may also contribute to mPFC-dependent behavior in the AST. We observed that restrained iAAV-Arc rats display increased time and number of trials required to reach criterion compared to restrained iAAV-scramble rats in the side reversal and light discrimination phases, which are regulated by the OFC and the mPFC, respectively (50). Impairments in the OFC-dependent side reversal phase suggest that some pPVT neurons may project to the OFC or to OFC-projecting mPFC neurons, and/or that the side reversal phase of the AST is not exclusively dependent on the OFC. It should be noted that other factors in addition to pPVT Arc, such as stress-induced noradrenergic neurotransmission in the mPFC (68, 69), may also contribute to improved cognitive flexibility following repeated restraint in male rats. To the best of our knowledge, our results are the first to identify a role of the pPVT in contributing to PFC-dependent cognitive flexibility.

Arc induction by stress and Arc actions are specific to the posterior division of the PVT. In the aPVT, restraint did not alter Arc expression or affect structural plasticity. This specificity is important because the PVT is an extensive midline thalamic nucleus with anterior and posterior divisions that differ significantly in their efferent projections. Whereas aPVT projections are widespread (21-23), projections from the pPVT are more exclusive and, in general, target stress-related brain regions like the mPFC and amygdala (17, 18, 20, 26). The differences in efferent projections from aPVT and pPVT neurons confer different functions and uniquely position the posterior division to regulate habituation to repeated stress, consistent with previous findings (14).

The findings presented here are the first to identify Arc as a key mediator of structural plasticity in the pPVT and for how individuals adapt to repeated stress. Much less is known about the function of Arc in the thalamus compared to other brain regions, like the hippocampus. Our results demonstrating may provide a foundation for understanding other plasticity-dependent processes regulated by thalamic circuits. Chemogenetic inhibition of the subpopulation of pPVT neurons projecting to the mPFC attenuated behavioral, but not neuroendocrine, habituation. This dichotomy suggests that pPVT projections to brain regions other than the mPFC (e.g. the BNST) may be responsible for pPVT regulation of the HPA axis. We also demonstrated that pPVT Arc is necessary for pPVT-mPFC coherence and PFC-dependent cognitive flexibility. Together, our findings provide novel insights into the molecular and network mechanisms underlying habituation to repeated stress, a fundamental and phylogenetically conserved property of the stress response.

## Acknowledgements

This work was supported by the Defense Advanced Research Projects Agency (DARPA) and the U.S. Army Research Office under grant number W911NF1010093 to SB. Additional support in the form of a Training Grant in Neurodevelopmental Disabilities NIH/NINDS T32 NS007413 and a NARSAD Young Investigator Grant from the Brain & Behavioral Research Foundation (29185) was awarded to BC. The authors have no conflicts of interest.

